# Modelling bacterial twitching in fluid flows: a CFD-DEM approach

**DOI:** 10.1101/648915

**Authors:** Pahala Gedara Jayathilake, Bowen Li, Paolo Zuliani, Tom Curtis, Jinju Chen

**Affiliations:** School of Engineering, Newcastle University, United Kingdom, NE17RU; School of Computing, Newcastle University, United Kingdom, NE17RU

**Keywords:** Bacterial motility, upstream twitching, modelling, CFD-DEM

## Abstract

Bacterial habitats are often associated with fluid flow environments. There is a lack of models of the twitching motility of bacteria in shear flows. In this work, a three-dimensional modelling approach of Computational Fluid Dynamics (CFD) coupled with the Discrete Element Method (DEM) is proposed to study bacterial twitching on flat and groove surfaces under shear flow conditions. Rod-shaped bacteria are modelled as groups of spherical particles and Type IV pili attached to bacteria are modelled as dynamic springs which can elongate, retract, attach and detach. The CFD-DEM model of rod-shape bacteria is validated against orbiting of immotile bacteria in shear flows. The effects of fluid flow rate and surface topography on twitching motility are studied. The model can successfully predict upstream twitching motility of rod-shaped bacteria in shear flows. Our model can predict that there would be an optimal range of wall shear stress in which bacterial upstream twitching is most efficient. The results also indicate that when bacteria twitch on groove surfaces, they are likely to accumulate around the downstream side of the groove walls.

## Introduction

A bacterial biofilm is a bacterial community attached into a surface through extracellular polymeric materials ^1^. Prior to biofilm formation, bacteria may need to deposit on the surface from their planktonic state. After bacteria deposit on surfaces they may “*twitch*” or crawl over the surface using appendages called type IV pili (TFP) ^2-5^ to “explore” the substratum to find suitable sites for growth and thus biofilm formation. Pili emanate from bacterial surface and they can be up to several μm long (though they are nm in diameter^6^). Bacterial twitching occurs through cycles of polymerization and de-polymerization of type IV pili ^7,8^. Polymerization causes the pilus to elongate and eventually attaching into surfaces. De-polymerization makes the pilus to retract and detaching from the surfaces. Pili retraction produces pulling forces on the bacterium, which will be pulled in the direction of the vector sum of the pili forces, resulting in a jerky movement (Figure 1). A typical TFP can produce a force exceeding 100 pN ^9^ and then a bundle of pili can produce pulling forces up to several nN ^10^. Bacteria may use pili not only for twitching but also for cell-cell interactions ^11,12^, surface sensing ^13,14^ and DNA uptake ^15^.

**Figure 1.**
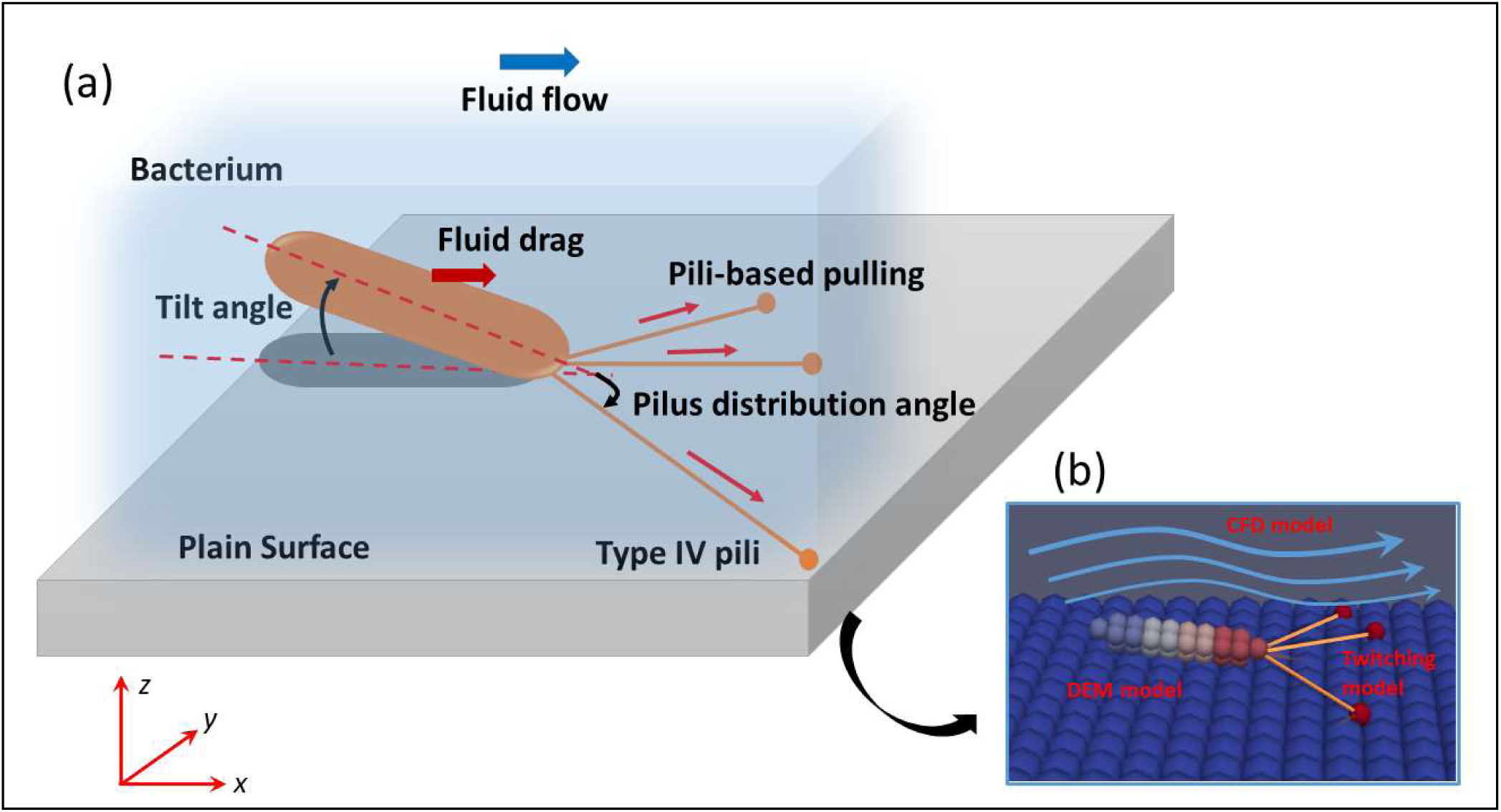
**(a)** Schematic of bacterial twitching; **(b)** CFD-DEM model. The rod-shaped bacteria are modelled as a group of spherical particles rigidly assembled together. The pili are emanated from the bacterial pole coloured in red. Each pilus is modelled as a dynamic spring which can elongate, attach, retract and detach from the surface.

Twitching motility could depend on many factors including surface properties, pili arrangement on bacterial surface, and environmental conditions such as oxygen concentration and fluid flow rate ^16^. For example, when pili emanate only at the poles of bacteria (e.g., *Pseudomonas aeruginosa*), the bacteria will have persistent motion ^17,18^. But, if pili are all around the cell body (e.g., *Neisseria gonorrhoeae*), the bacteria will have trapped or diffusive motion due to the *tug of war* mechanism ^19,20^. If a pilus detaches while all the pili are in high tension and anti-parallel configuration, the bacterium will suddenly align along the resultant direction of the remaining bounded-pili causing a sudden change of the twitching direction. This is the so called *slingshot motion* and bacteria may use this mechanism to change crawling direction ^3,4^. Bacterial twitching will depend on some physicochemical and structural properties of the surface. For instance, the pili attachment is enhanced ^2,18,21^when the substratum is covered by extracellular polymeric materials. Patterned surfaces can be a barrier for bacterial twitching and hence hinder surface exploration by bacteria ^7,22^. Chang, et al. ^22^ have shown that micro-scale surface topography (pillars) appears to be a barrier to the surface motility of *Pseudomonas aeruginosa* and it may hinder the ability of such cells to explore a surface. However, when the surface has micro-scale grooves, bacteria may display persistent twitching along grooves because cells can be guided by the groove walls ^2,23^. Bacteria can also differently deploy pili ^17^ and change pili retraction speed ^24^ to adapt to nutrient availability. In fluid flow environments, rod-shaped bacteria tend to twitch against the flow because the fluid flow tends to align the bacteria along the flow direction while they are anchored to the tethering points, and then the fluid drag causes bacteria to flip around the anchoring point and twitch upstream (see Figure 1) ^25-27^.

The experimental visualization of pili is difficult requiring great skill and specialised equipment ^28^. Therefore, mathematical modelling of TFP mediated bacterial twitching is vital to understand the twitching mechanism under different environmental conditions. Researchers have already modelled twitching motility of bacteria using a variety of mathematical models. For instance, Marathe, et al. ^20^ modelled *Neisseria gonorrhoeae* as point particles and used stochastic pili dynamics to simulate a *tug of war* mechanism with directional memory of twitching action. This work reported that directional memory enhances the surface exploration of bacteria. Molecular dynamics (MD) or discrete element based methods (DEM) have been widely used to understand bacterial twitching. Brill-Karniely, et al. ^4^ used a kinetic Monte Carlo algorithm together with MD to model TFP mediated twitching of *Pseudomonas aeruginosa*. This work reported that a minimal amount of angular rigidity of pili is needed to produce some experimentally observed behaviours of twitching bacteria. Furthermore, this work revealed that two TFP can produce the recently observed *slingshot motion* ^3^ when one pilus releases at a high-tension anti-parallel configuration of two pili. More MD based twitching models include de Haan ^6,^Zaburdaev, et al. ^19,^Ryota Morikawa ^29^. However, these very interesting models have not considered interactions of a twitching bacterium and its hydrodynamic environment. This represents an important gap in our knowledge because the hydrodynamic environments can completely change twitching direction (e.g., upstream twitching) as well as influencing deposition and detachment ^30^. In the present work, three-dimensional Computational Fluid Dynamics coupled with Discrete Element Method (CFD-DEM) is used to model rod-shaped bacterial twitching on flat and groove surfaces under various shear flow conditions. Various forms of CFD-DEM models have been employed to study bacterial deposition before ^31-33^. The novelty of our model is the use of a three-way coupled (two-way coupled fluid-cell interactions plus cell-cell interactions) CFD-DEM model together with pili dynamics to study bacterial twitching on flat and groove surfaces with fluid flowing over the surfaces. The model is implemented on an open source CFD-DEM package called SediFoam ^34^. The method is used to predict some experimentally observed behaviours of bacteria twitching in shear flows such as upstream twitching ^26^. The model is generic in nature, but the parameters are chosen such that they are relevant to the *Pseudomonas aeruginosa*.

## Results and Discussion

When immotile rod-shaped bacteria move in shear flows they will freely orbit in shear flows ^30^ which is called “*Jeffery orbiting*”. We first compare the orbiting of a rod-shaped bacterium with theoretical results to validate the CFD-DEM model. Then, the model is used to study bacteria twitching on a rough surface in the presence of a static fluid medium. Finally, the model is employed to investigate bacteria twitching in a flowing environment on a rough-flat and rough-groove surface.

The computational domain for the following simulations is a rectangular box having the dimensions of [0, 50] × [0, 20] × [0, 20] μm^3^. Periodic velocity boundary conditions in two horizontal directions (*x* and *y* direction) and no-slip and fixed-velocity boundary conditions are applied respectively at the bottom and top walls (*z* direction). Pressure is periodic in the horizontal directions and zero gradient boundary conditions are applied at the top and bottom walls. The parameters used for the following simulations are listed in Table S1. A single bacterium is simulated unless specified otherwise and the bacterium is initially oriented in the flow direction.

### Model validation for *Jeffery Orbiting*

SediFoam has been extensively validated for spherical particle laden flows ^34-36^. We use SediFoam for rod-shaped objects in this work and hence we validate the model for *Jeffery orbit* before using for bacterial twitching. The analytical expression for orbiting angular velocity 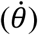 and period (*T*) of a rod-shaped bacterium having an aspect ratio of *a* in a shear rate of 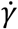 are given by Jeffery ^37^ as

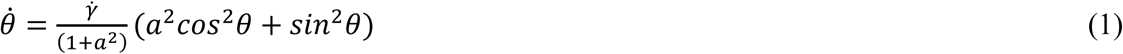

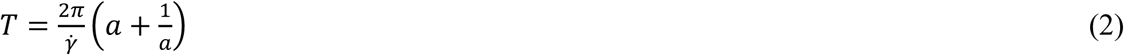

The CFD-DEM model is validated for the *Jeffery orbit* at different shear rates and aspect ratio of the cell body. The analytical solution for the orbiting angular velocity and the period of the orbit are compared with the present numerical results. Figure S1 (a) shows the numerical and analytical results at *a* = 3 and 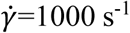 and it can be seen that the present CFD-DEM model can predict the orbit transit of a rod-shaped bacterium in shear flows accurately. Figure S1 (b) compares the periods at different aspect ratios and shear rates and it is evident that the model is capable of predicting the theoretical results. The relative error of the maximum and minimum angular velocities are presented in Figure S1 (c) and it can be seen that the relative error is less than 15% for all the cases we have considered here. A relative error as large as 15% would be because the analytical solution is valid only for inertialess rods, the present CFD-DEM model computes only average hydrodynamics around the bacterium, and the shape of the bacterium is not precisely a rod. Therefore, a relative error of 15% would be still acceptable for reasonable predictions of rod-shaped bacteria interaction with fluid flows.

### Bacterial twitching in static fluid

Bacterial motility would be affected by the number of pili and how those pili distribute at the bacteria poles ^2,17^. Therefore, the present model is employed to understand how the number of pili and the distribution angle (*α*) influence twitching characteristics. Bacterial twitching is usually characterised by the Mean Square Displacement (MSD) which can explain the twitching behaviour (diffusive, trapped, and persistent) based on the MSD power (MSD = Kt^n^, where n is the MSD power, K is a constant, and t is time) ^4,20^. Figure 2 shows the MSD power for different pili distribution angles and pili numbers. As the pili angle increases the MSD power decreases because the cell is more likely to trap between pili which are in force equilibrium. The numbers shown in bars are the R^2^ value of the regression to compute MSD power and it can be seen that it decreases as the pili distribution angle increases, because the cell has more irregular motion in that case. When the number of pili increases at the same pili angle the MSD power does not change much. The trajectory of the leading and trailing poles are shown in Figure 2(b) when the pili number is 2 and the pili distribution angle is 30^0^. The trailing pole moves above the leading pole because of the inclination of the bacterium to the surface, as observed experimentally by others ^17^.

**Figure 2.**
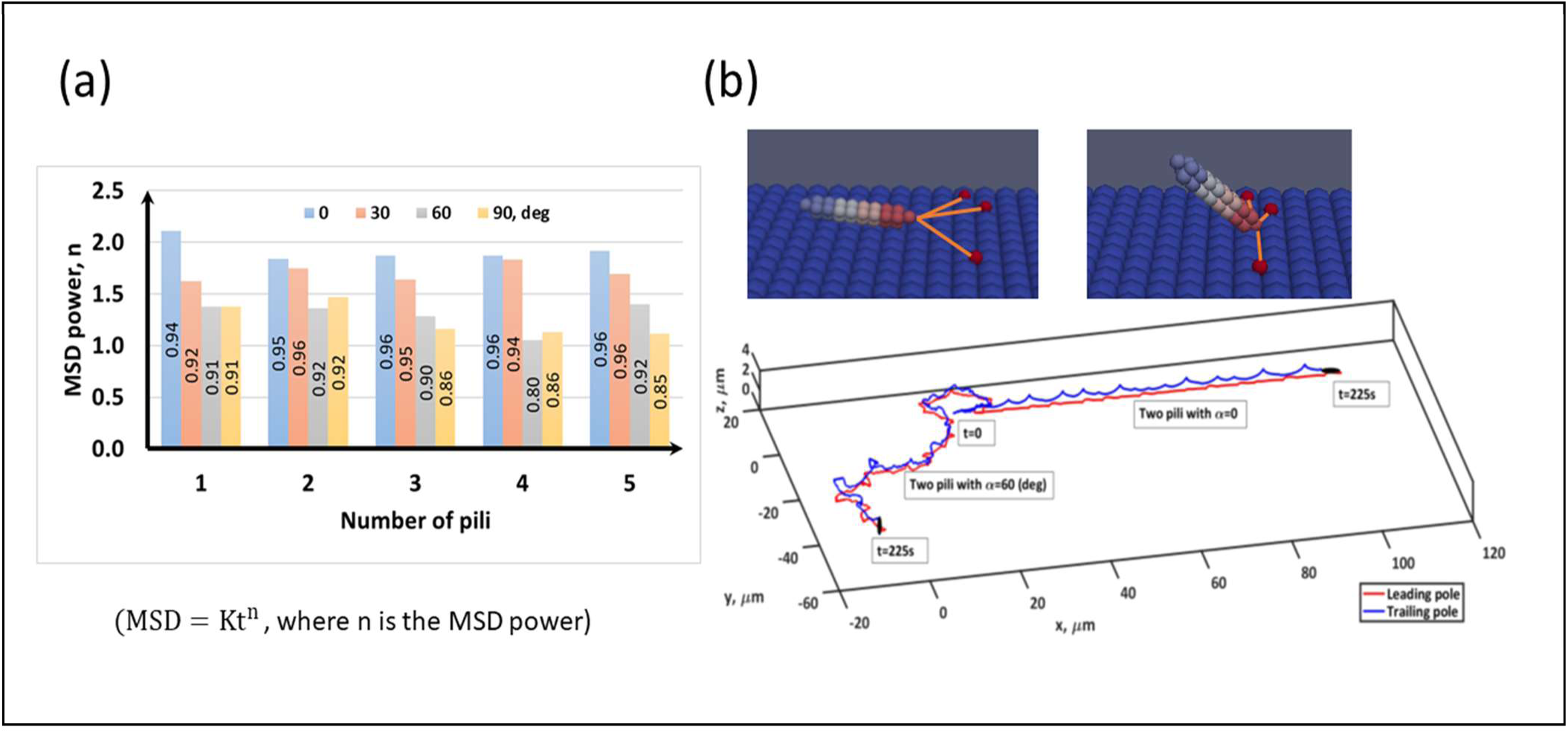
**(a)** MSD power for different pili angles and pili numbers (MSD = Kt^n^, where n is the MSD power, K is a constant, and t is the time); **(b)** When the bacterium is pulled by the pili for an extended period of time, the bacterium gradually gets inclined to the surface and if the period is long enough the bacterium would reach to a vertical orientation. The trajectory of the leading and trailing poles are shown for pili number is 2 and the pili angle is 30 deg.

Figure 3 shows the variation of twitching velocity (*V*_*t*_) for different pili distribution angles and pili numbers. As expected, the quasi-stationary time (time spent at *V*_*t*_ <0.01 μm/s) decreases and moving time (time spent at *V*_*t*_ >0.01 μm/s) increases when the number of pili increases, since the bacterium is pulled by pili more frequently. The pili distribution angle has significant influence on the intermediate velocity (0.01< *V*_*t*_ <0.8 μm/s). The average twitching velocity is less than 0.8 μm/s for all the cases (Figure 3b) which is a realistic prediction for the twitching velocity of *Pseudomonas aeruginosa* (i.e. 0.3μm/s) found in Maier and Wong ^2,^Jin, et al. ^3^. The average twitching velocity is more sensitive to the number of pili than the pili distribution angle. These results indicate that the MSD of bacteria can be simply written as MSD = K(N_p_)t^n(α)^ because the MSD power and the twitching velocity are more sensitive to the pili distribution angle and the numbers of pili, respectively. Here, *n*(*α*) is the MSD power and it explains the nature of the twitching motility, which is sub-diffusive (trapped) when *n*(*α*) <*1* and super-diffusive (persistent) when *1* < *n*(*α*) < *2*, diffusive when *n*(*α*) =1 and ballistic when *n*(*α*) = 2. In the present study, it can be seen that the twitching motility is super-diffusive most of the time (Figure 2a) and it never has a sub-diffusive motion. This is expected because all the pili are focused at one pole and their distribution angle is also taken from the Normal distribution and hence bacteria have persistent motion. There is experimental evidence that *Pseudomonas aeruginosa* would twitch with a MSD power of 1.55±0.34 when they twitch using unipolar TFP ^17^.

**Figure 3.**
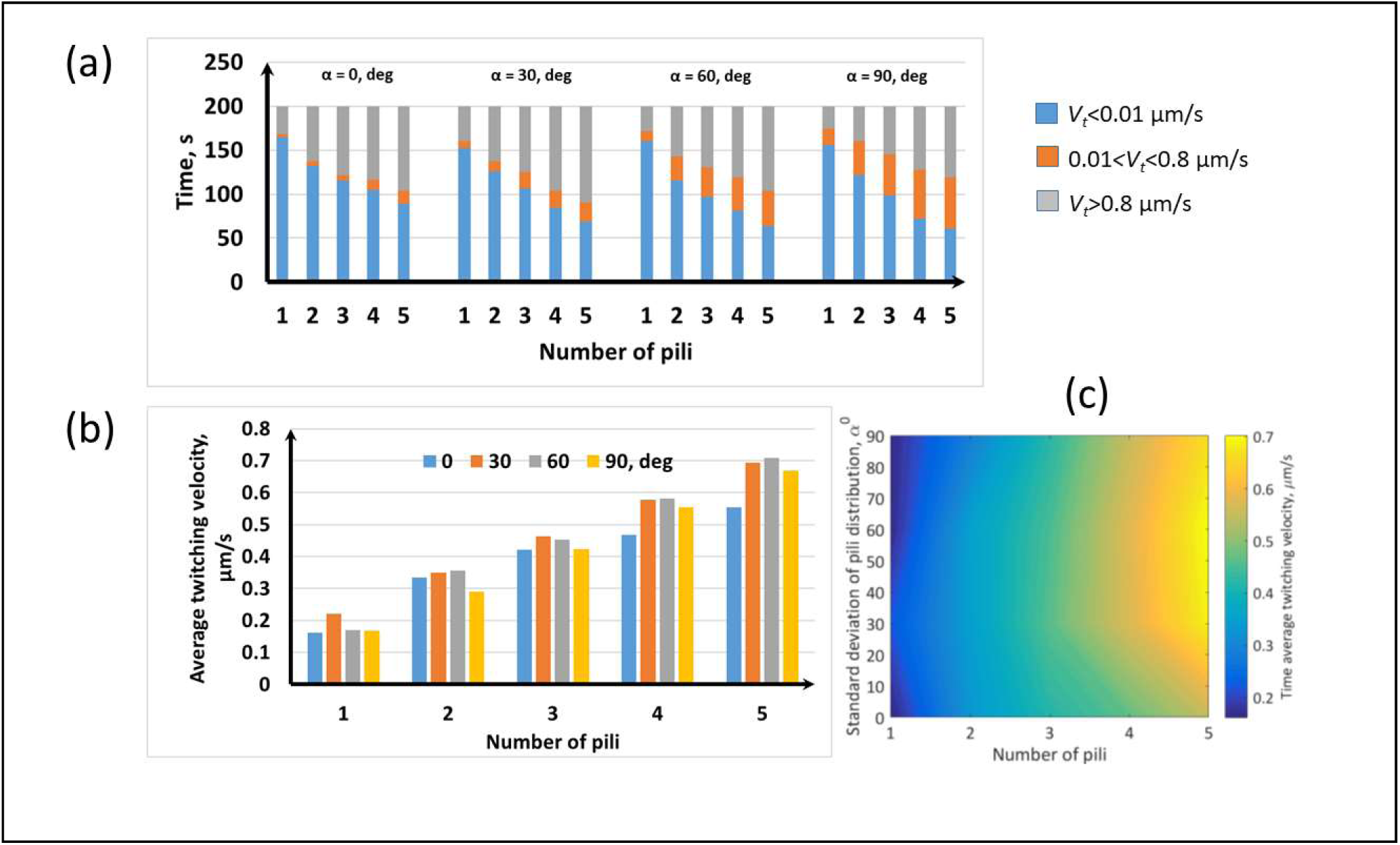
**(a)** Twitching velocity for different pili distribution angles and pili numbers. As expected, the stationary time (*V*_*t*_<0.01μm/s) decreases and moving time (*V*_*t*_ >0.8μm/s) increases as the number of pili increases because the bacterium is pulled by pili more frequently then. Pili distribution angle has significant influence on the intermediate velocity (0.01< *V*_*t*_ <0.8μm/s); **(b-c)** The average twitching velocity is more sensitive to the number of pili than the pili distribution angle.

Figure 4 shows the tilt angle for different pili distribution angles and pili numbers. With increasing number of pili the tilt angle distributes in a wide range, while the average tilt angle is still around 5 to10 degrees. The twitching experimental data of *Pseudomonas aeruginosa* reported in Ni, et al. ^17,38^ indicated that the average tilt angle would be around 15 degrees and our results are in a reasonable range considering the assumptions of the model. Our results show that as the pili distribution angle and numbers of pili increase there is a tendency for the bacterium to trap in a vertically-oriented configuration (Figure 4c-d). This is rather similar to the vertically-oriented upright walking of *Pseudomonas aeruginosa* ^17,38,39^. The model shows that when a cell is trapped between pili for an extended period of time, the cell has more time to rotate around its body and reach a vertical orientation. However, our model is not capable of capturing the upright walking motility of bacteria. In the present model, vertically oriented bacteria remain trapped and then gradually move to the horizontally-oriented configuration and crawl when the trapped-configuration of pili is changed once a new pilus attaches or breaks, and the force becomes unbalanced. It appears that a special pili dynamics mechanism will be needed to capture those vertically-oriented upright walking bacteria and that is out of the scope of this paper.

**Figure 4.**
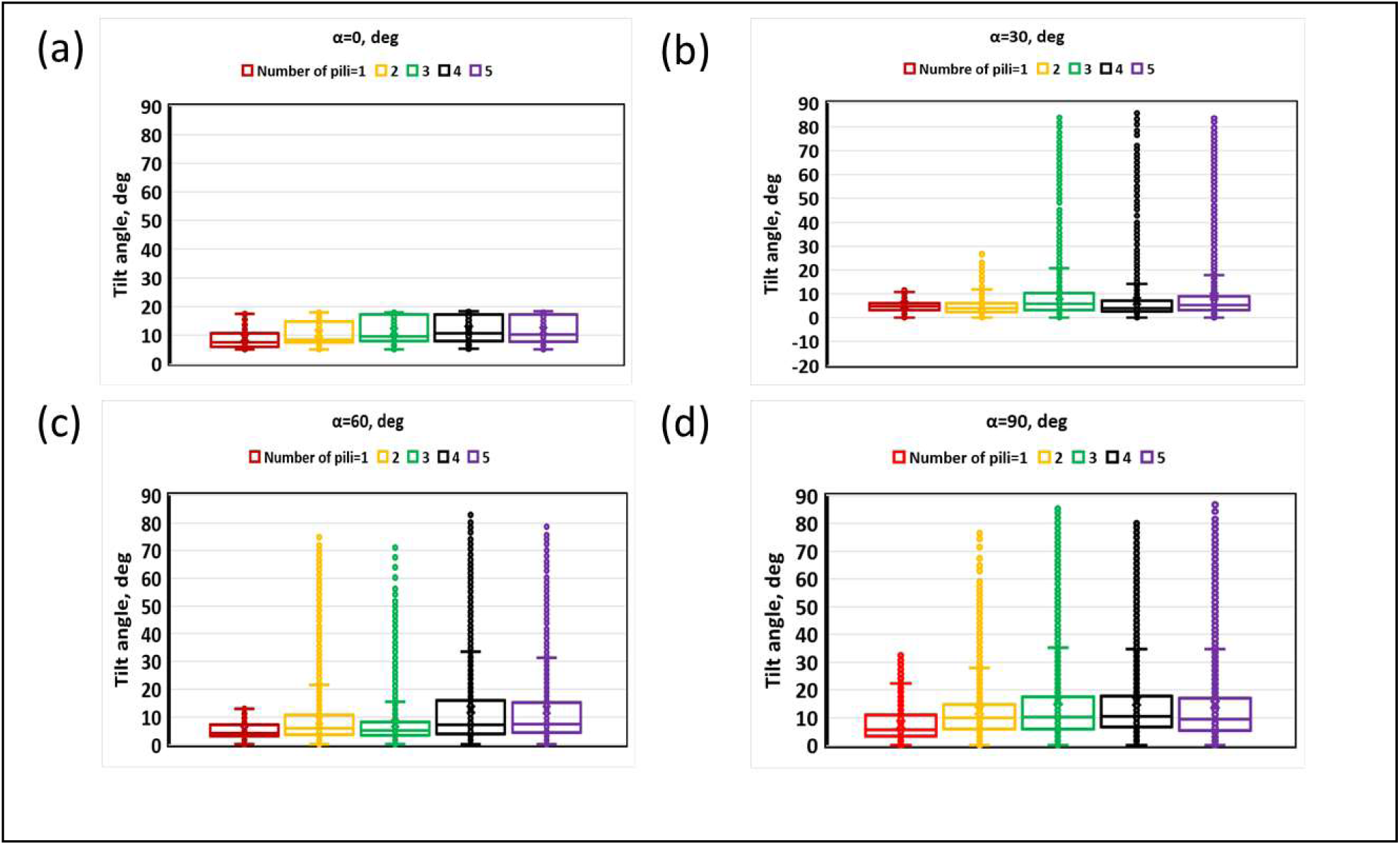
Tilt angle for different pili distribution angles and pili numbers: **(a)** 0; **(b)** 30; **(c)** 60; **(d)** 90 degrees. Larger the number of pili the tilt angle distributes in a wide range while the average tilt angle is around 5-10 degrees. When the pili distribution angle and the number of

### Bacterial twitching in flowing fluid

We study bacterial twitching under a range of flow velocity (0-4 mm/s) which corresponds to a range of wall shear stress values (wall shear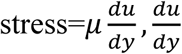 is the shear rate at the wall). We study a bacterium having two pili with 30^0^ distribution angle and with increased pili elongation velocity (10 times) for the following reasons. When we add many pili a smaller time step is needed for scaling up (≪0.1 s) to maintain numerical stability which has a significant computational cost when flow fields are taken into account. Even if we use two pili for the model, it does not necessarily mean that the bacterium has only two pili. Because of increased elongation velocity these two pili can mimic several pili in a real system because a new pilus is created faster after the breakage of an existing pilus. Figure S2 shows the main events associated with bacterial twitching in a flow environment, which are upstream twitching, detachment, orbiting in shear flow and re-attachment.

Figure 5 shows the probability of direction of motion of the cell at different wall shear stresses. If the fluid is static (Figure 5a), the cell can twitch in any direction on the surface because there is no preferential driving force, and therefore the probability of twitching direction being in each angular bin is about 1/12=0.08. Then, when the fluid flows, the cell tends to twitch upstream as seen in Figure 5(b-d) indicating the increased probability of bins from 90^0^ to 270^0^ compared to the no-flow scenario. For the selected wall stresses, the maximum upstream twitching probability occurs at around 0.1 Pa and that probability decreases as the wall stress is either increased or decreased from that value, indicating that there would be an optimal flow condition for bacterial upstream twitching. Figure 6(a) supports this finding and shows a sinusoidal variation of twitching probability with wall shear stress. The probability of upstream twitching decreases and reaches a minimum and then increases to a maximum and then it decreases again. The reason for this behaviour is that the cell is initially headed in the flow direction and at low shear stresses the bacterial cell is not subjected to sufficient shear forces to rotate it in the upstream direction. Therefore, the cell will actively twitch and be passively advected in the flow direction. But, at moderate shear stress, the cell will be rotated and faced in the upstream direction and it will then twitch against the flow. As the fluid flow further increases, the fluid drag forces would tend to dominate over to the pili-based pulling forces and hence upstream twitching decreases again. Figure 6(b) shows the time average velocity in upstream and downstream directions. Upstream twitching velocity is fairly constant for a range of shear stress, in agreement with experimental findings in the literature ^26^. It can be seen that the upstream twitching distance has a unimodal distribution (Figure 6c) with wall shear stress. The fluid flow conditions, apart from the optimal wall shear stress (that is around 0.1 Pa), may adversely influence upstream twitching. Shen, et al. ^26^ showed that upstream twitching of *P. aeruginosa* would be most efficient when the wall shear stress is around 0.5 Pa, and our model predictions are also in the same order. It can be seen in Figure 6(d) that bacteria detach from the wall more frequently when the wall shear stress is more than the optimal stress. This is because the fluid drag is dominant to the pili-based pulling when the stress is far beyond the optimal value.

**Figure 5.**
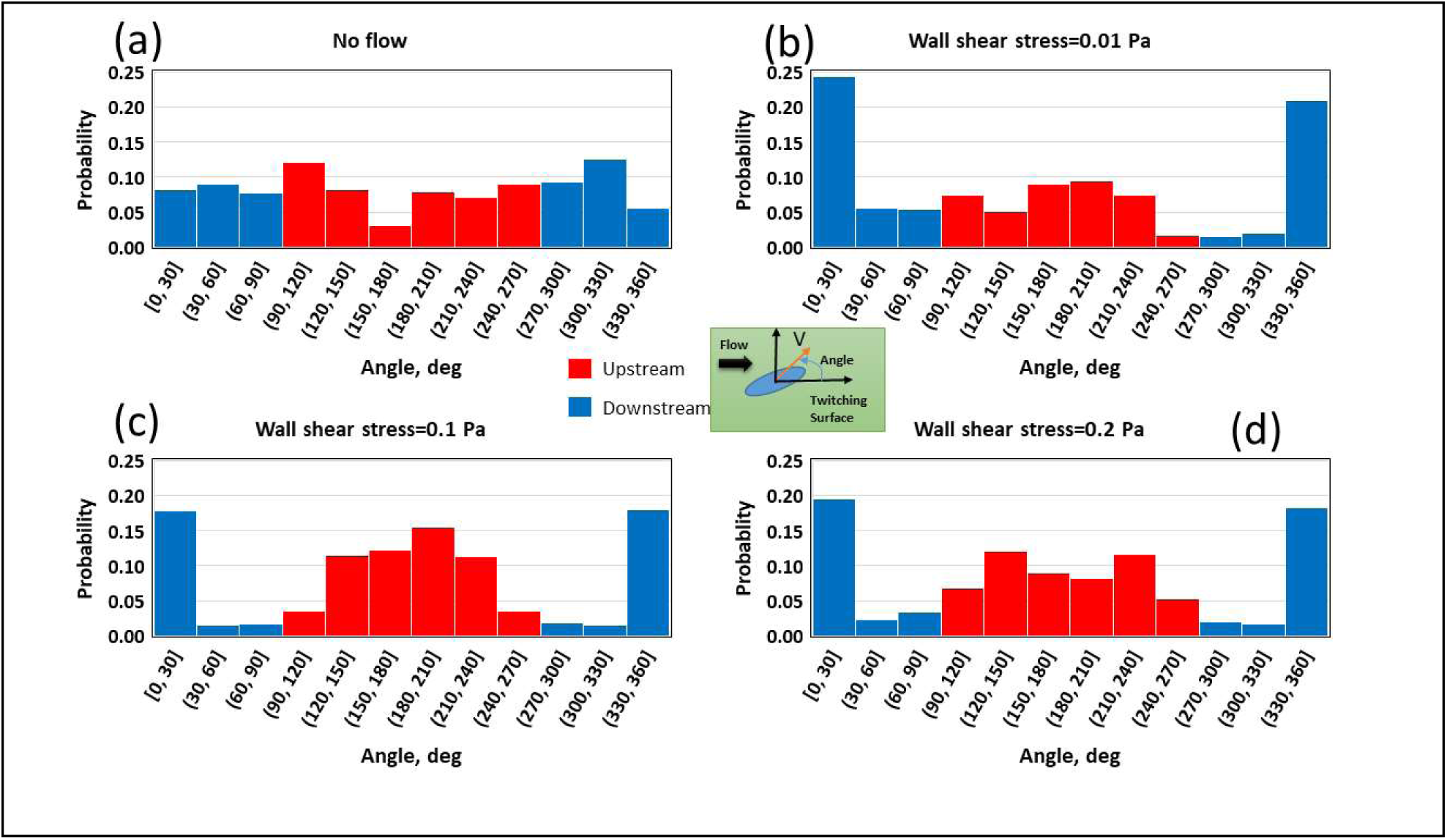
**(a)** Probability of the direction of motion of the cell is shown as a function of wall shear stress. If the fluid does not flow, the cell can twitch any direction on the surface (the probability of each angular bin is around 1/12=0.08; **(b-d**) As the fluid flow (or wall shear stress) increases, the cell tends to move upstream, the probability of bins from 90 to 270 degrees increases.

**Figure 6.**
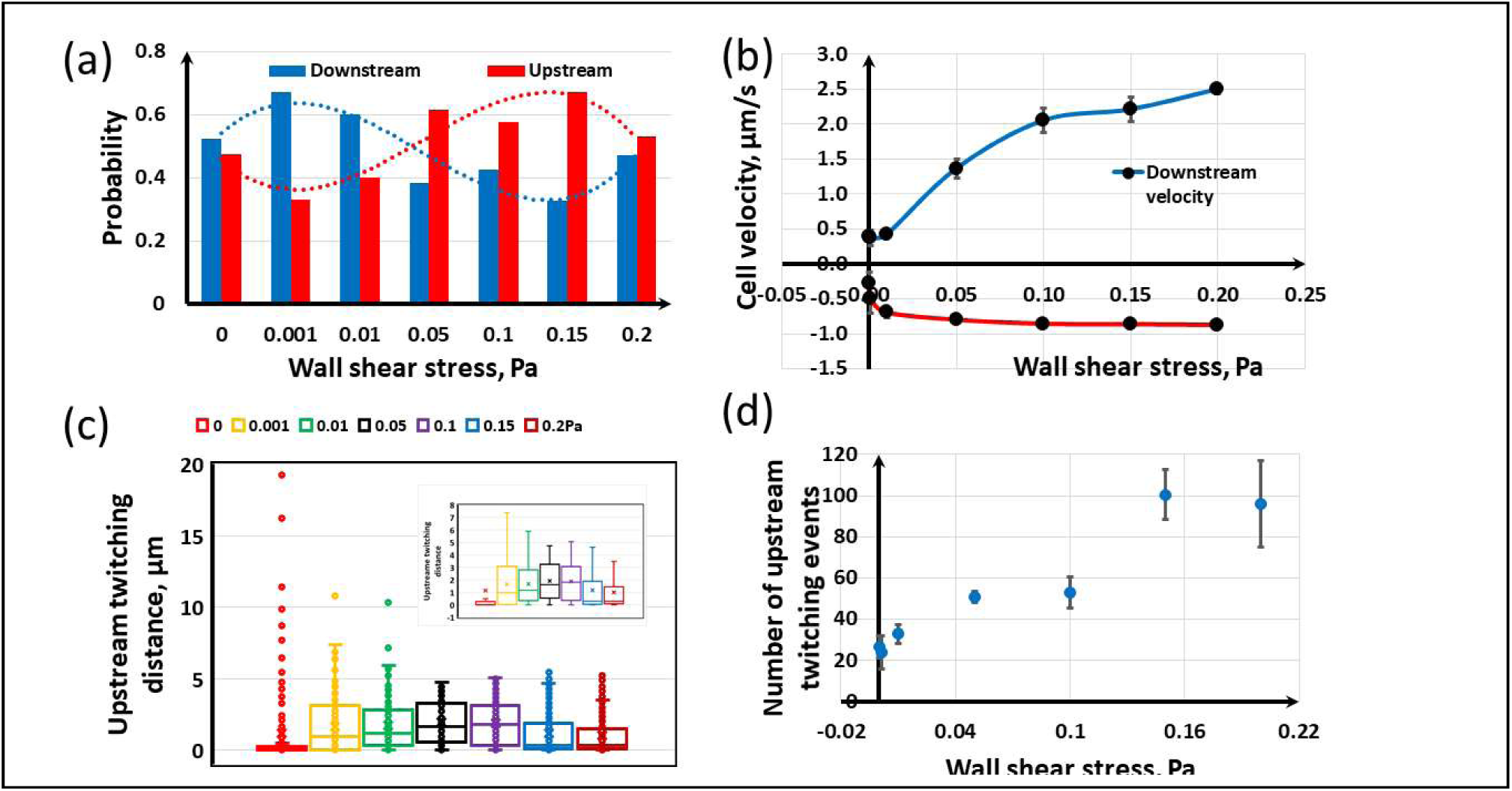
**(a)** As the fluid flow increases, the probability of upstream twitching (Red bars) decreases and reaches a minimum and then increases to a maximum and then it decreases again; **(b)** Time average velocity in upstream and downstream directions; **(c)** Upstream twitching distance; **(d)** Number of twitching events.

### Bacterial twitching on a groove surface with fluid flow

It is important to study how imposed fluid flows would influence bacterial twitching on structured surfaces because twitching would be influenced by both structures and moving fluid in this case. Therefore, we study bacterial twitching on a groove surface (Figure S3) when fluid flows across the grooves. The cross section of each groove wall (protrusion) is in the order of bacterial size and it is chosen as 6×5 μm^2^ (wide ×height). Bacterial upstream twitching is investigated at two different groove widths (19 and 46 μm, which are about two and four times of the bacterial length, cell body plus pili length) at a wall shear stress of 0.15 Pa. A similar geometry has been experimentally investigated for *Escherichia coli* deposition in Gu, et al. ^40^. The pili dynamics is similar to the previous case of bacterial twitching on flat surface under shear flows (i.e., two pili with distribution angle of 30^0^). Four bacteria are randomly seeded on the surface. Figure S3 shows bacterial twitching on flat and groove surfaces. As expected, bacteria are trapped and twitch along the grooves for the non-flat surfaces.

Figure 7 shows the probability of twitching direction at different surface conditions. As also shown in Figures 5-6, it can be seen that bacteria simply twitch upstream on the flat surface (Figure 7a). Figures 7 (b, c) indicate that the groove width has a vital influence on twitching motility. Upstream twitching is inefficient for the narrow groove (Figure 7c) and the direction of motility is chaotic in that case. The reason is that bacteria frequently collide on the groove wall because of fluid drag and upstream twitching resulting in the direction of motility change regularly. The groove walls also guide bacteria to twitch along the grooves. These constraints would give bacteria uniform chances to move in any direction on the groove when the groove width is relatively low. Figures S3 and 8 indicate that bacteria tend to accumulate downstream of the groove walls. This phenomenon would be theoretically meaningful because the fluid drag behind the walls would be weak and therefore bacteria would not be easily pulled along the flow. Therefore, bacteria that twitch upstream and reach the walls are likely to reside there for an extended period.

**Figure 7.**
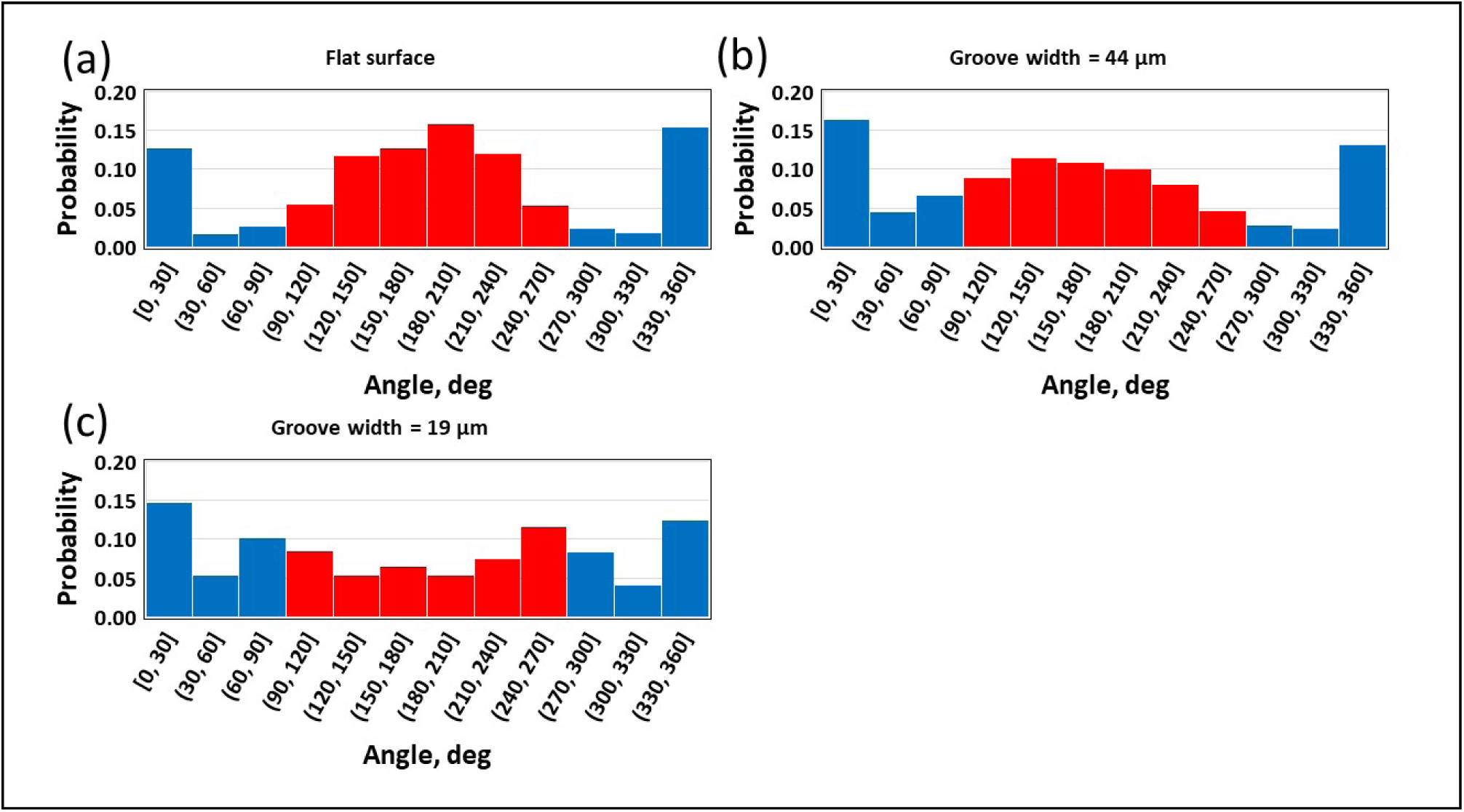
Effect of surface topography on bacterial upstream twitching at wall shear stress of 0.15 Pa: **(a)** Flat surface; **(b)** Groove width is 44 μm; **(c)** Groove width is 19 μm. It is seen that the bacteria have a chaotic motion when the groove width is smaller.

**Figure 8.**
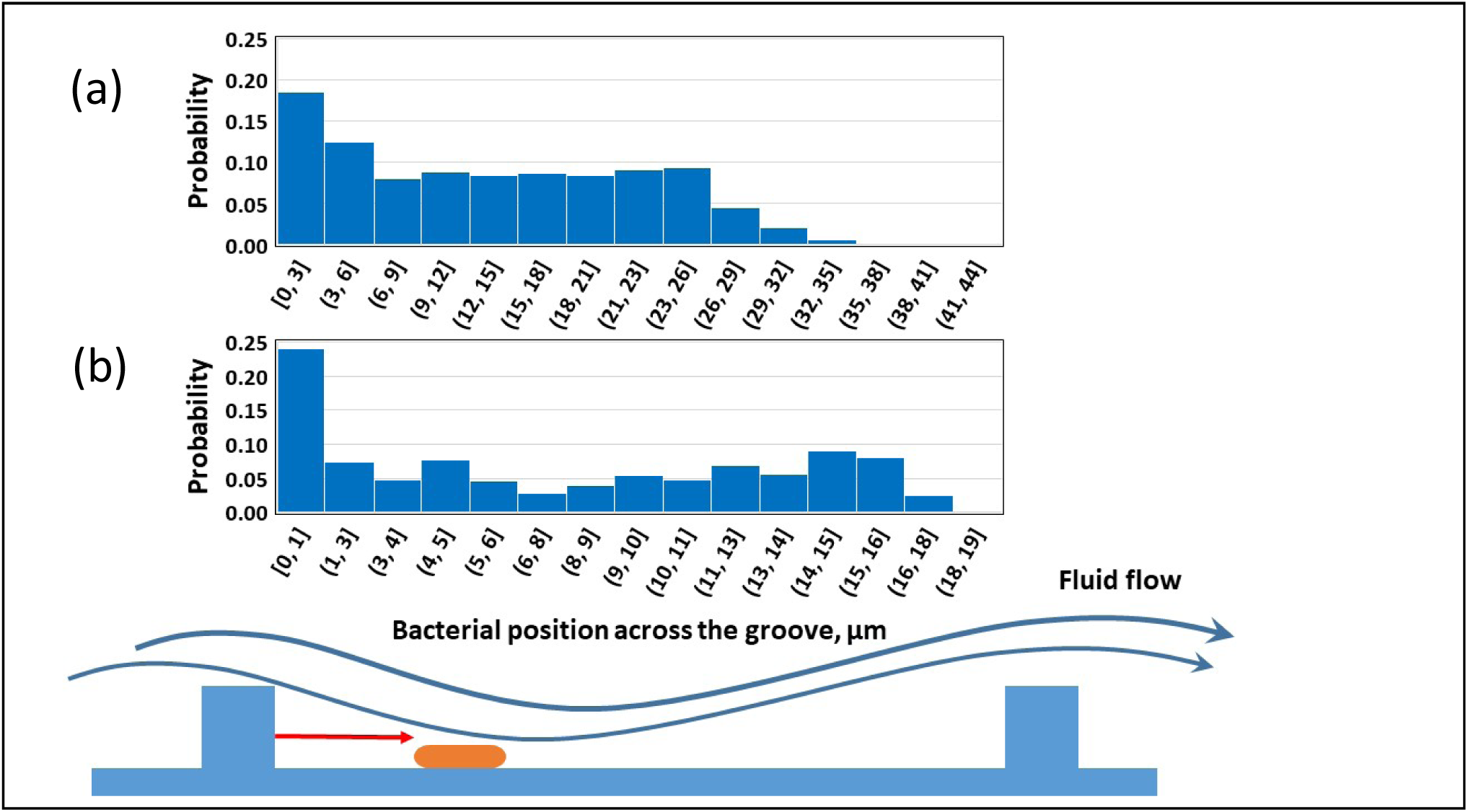
Probability of bacterial residency time at difference places inside the groove: **(a)** Groove width is 44μm; **(b)** Groove width is 19μm. The results indicate that bacteria are likely to accumulate near downstream sides of groove walls.

## Conclusion

Bacterial motility shows interesting phenomena when active motility (swimming, twitching and so on) interferes with a surrounding fluid ^26,30,41^. Upstream twitching is a mechanism used by rod-shaped bacteria such as *Pseudomonas aeruginosa* to colonize upstream sites of flow environments such as catheters. In this work, a CFD-DEM model is used to study bacterial twitching in fluid flows. The model can predict super diffusive motility in static fluid, upstream twitching in fluid flows, and flow-induced cell detachment/re-attachment to the surface. In agreement with experimental findings of Shen, et al. ^26^, our model can predict that there would be an optimal range of wall shear stress in which bacterial upstream twitching is most efficient. When bacteria twitch on a groove surface, the resultant effect of fluid flow and surface topography would decide the nature of twitching and spatial segregation of bacteria on the surface. While our model can predict general characteristics of bacterial twitching, the model should be carefully validated against experimental data before it can be used to gain more detailed insights about bacterial twitching in fluid flows.

Even though our model was basically used to study upstream twitching, it can give some insights for variety of other twitching phenomena. The present model can be used to investigate bacterial twitching on compliant surfaces and other surfaces with complex micro or nano scale structures. The model can be a robust tool to study twitching motility of different shapes of bacteria (e.g., *Neisseria gonorrhoeae* and *Synechocystis sp PCC* 6803) ^2^ and TFP-based colonization of curved shape bacterium *Caulobacter crescentus* in fluid flows ^42^. Moreover, our model could be used for investigating how the oscillatory localization of TFP (dependent on nutrient conditions) of *Pseudomonas aeruginosa* and *Myxococcus xanthus* would interfere with fluid flows^17^.

## Methodology

We have implemented twitching dynamics of bacteria into the existing CFD-DEM platform called SediFOAM ^34^, which couples the molecular dynamic code LAMMPS ^43^ and the well-established CFD package, OpenFOAM ^44^. The present work is an extension for the authors’ Individual-based model of microbial communities implemented on LAMMPS^45^. SediFOAM has been primarily used to simulate particle sedimentation in fluid. In the present work, we extend SediFOAM to model rod-shaped bacteria twitching in fluid flows. The model components are explained below.

### Discrete element modelling (DEM) of bacteria and surface

We model bacteria and solid substratum (flat and groove surfaces) by using spherical particles. Rod-shape bacteria are modelled as a rigid assemble of several spherical particles (See Figure 1). The total force on the rigid body is computed as the sum of the forces on its constituent particles. This idea has been employed before for modelling rod-shaped bacteria ^33^. The translational and rotational movement of the rigid body is calculated based on Newton’s second law as

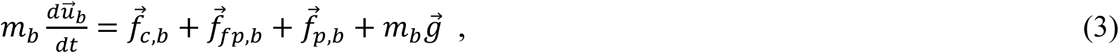

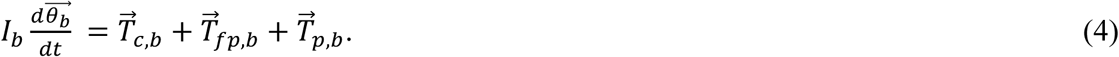

Here *m*_*b*_ and *I*_*b*_ are the mass and moment of inertia of the bacterium (rigid body), respectively. Eq (3) describes the translational velocity 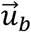 of the bacterium, and the four terms on the right hand side represent respectively the contact, fluid interaction, TFP pili, and gravitational forces acting on the cell. The rotational movement of the cell body 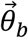 is calculated based on the torque produced by contact forces 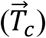, fluid interaction forces 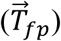 and pili forces 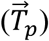. The contact forces are calculated based on Hook’s law depending on the overlap distance between interacting particles. Fluid interaction and pili forces are further explained below.

### Computational fluid dynamics (CFD)

The fluid is assumed Newtonian and its flow is described by the locally-average Navier-Stokes equation as

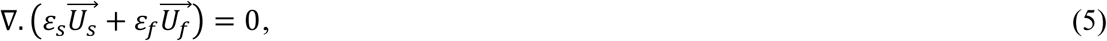

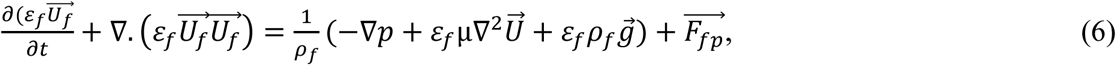

where *ε*_*s*_ is the solid volume fraction and *ε*_*f*_ =1 − *ε*_*s*_ is the fluid volume fraction. The fluid density *ρ*_*f*_ and its viscosity µ are assumed as constants. Here 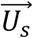 and 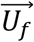 are the velocity of the solid and fluid phases, respectively. The gravity 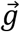 is also included because fluid and bacterial density would be different and hence buoyancy forces would be important. The last term 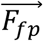 represents fluid-solid interaction forces, which are drag, lift, added mass, and lubrication forces as detailed in the SediFOAM documentation ^34^ and not repeated here. The Eulerian fields *ε*_*s*_, 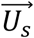 and 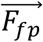 are calculated by averaging the information of Lagrangian particles.

### Twitching model

The TFP are modelled as dynamic springs emanating from one pole of the bacterium and these springs can elongate, retract, attach and detach from the surface (Figure 1). Each pilus operates independently from the others. When a new pilus is born at the bacterial pole, its angular direction from the bacterial axis is randomly decided according to a Normal distribution with standard angular deviation of *α, N*(0, *α*^*2*^). After the pilus elongates at constant velocity *ν*_*e*_ to a maximum length *L*_*max*_, it will attach to the surface with probability *p*_*a*_ and then it immediately starts to retract at a variable retraction velocity. A bound retracting pilus can detach (or break) with probability *p*_*b*_. Each un-attached or broken pilus retracts at velocity *ν*_*r*_ to the pole until it disappears and then a new pilus is born at the same pole at a random direction chosen from the Normal distribution. The total numbers of pili remain constant at any given time.

The pulling force of each pilus is modelled by assuming a linear spring with variable equilibrium length as

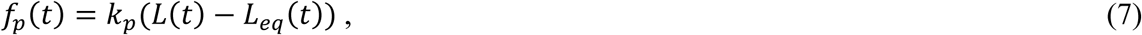

where *L*(*t*) and *L*_*eq*_(*t*) are the total length and equilibrium lengths of the pilus, respectively. The total length is simply the distance between the bacterial pole and the pilus tip. If the pilus is unbounded the equilibrium length is equal to the total length, which means the pulling force is zero. Once the pilus attaches to a surface the equilibrium length decreases representing pili retraction. As the retraction velocity of bounded pili depends on the pulling force ^7^, the equilibrium length is decreased as

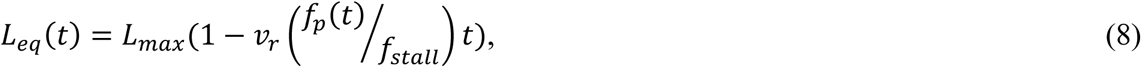

where *f*_*stall*_ is the maximum pulling force which can be produced by each pilus. A bounded pilus will break in a time interval Δ*t* with probability 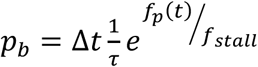 ^11^where *τ* is the characteristic time of pili detachment.

### Scaling-up

The time scale for twitching is much larger than 1s ^2,8^ and the time scale for fluid flow is much smaller than 1s. Therefore, twitching dynamics and fluid flow occur at two different time scales and hence it is needed to separate these time scales for the model. The CFD-DEM is run until quasi-steady state and the steady state flow field is calculated for a given bacterial position and orientation (Eqs. 3-6). Then, the bacterial twitching dynamics is calculated with a larger time step (Eqs. 7-8) and the position and orientation of the bacterium are updated using the velocities calculated from the CFD-DEM. Pili detachment and attachment events are also updated during this step. Next, the flow field is updated according to the new cell position and orientation through CFD-DEM calculations and so on. Therefore, three different time steps are involved in this model: the smallest time step of 10^−9^ s for DEM (Eqs. 3-4), an intermediate time step of 10^−5^ s for CFD, and the largest time step of 0.1s for pili dynamics (elongation, retraction, attachment, detachment) and scaling-up.

### Implementation in SediFoam

We have implemented our rod-shape bacterial model in SediFoam, in particular, in its LAMMPS module. Rod-shaped bacteria are created by assembling spherical particles rigidly by using the constraint *fix rigid* command provided by LAMMPS. Pili emanate from one pole of the bacterium. The *fix spring* command of LAMMPS is modified to model dynamic springs for TFP. Rough and irregular substratum is created using spherical stationary particles using the *fix move* command of LAMMPS (see the LAMMPS documentation at http://lammps.sandia.gov). Then, DEM is resolved by using the Verlet algorithm in LAMMPS; the PISO algorithm is used for solving CFD in OpenFOAM; and SediFOAM acts as an interface to transfer and map the properties of the Eulerian mesh and Lagrangian particles between the two modules.

## Data Availability

The datasets generated and analysed during the current study are available from the corresponding authors on request.

## Acknowledgement

The authors would like to thank the UK Engineering and Physical Sciences Research Council (EPSRC) for funding this work as part of the Newcastle University Frontiers in Engineering Biology (NUFEB) project (EP/K039083/1). This work would not have been possible without the multi-disciplinary collaboration enabled by this award.

## Author Contributions

All authors contributed to this work. PGJ, BL and JC designed the research. PGJ and BL developed the code. PGJ performed all the simulations and produced the data. PGJ and JC did data analysis and prepared the original draft. BL, PZ and TC reviewed and edited the manuscript. All authors have given approval to the final version of the manuscript.

