## Supporting Material for "Modelling bacterial twitching in fluid flows: a CFD-DEM approach"

**Table S1.** Parameters used for the simulations.

| Parameter | Value | Ref. |
|---|---|---|
| Flow chamber | 50×20×20 $\mu m^3$ | - |
| Fluid shear rate | 0-200 $s^{-1}$ | (1) |
| Fluid Kinetic viscosity | $1\times10^{-6}$ $m^2/s$ (*water*) | - |
| Bacteria size | 5×1 µm, length × diameter | (2) |
| Bacterial and surface stiffness | $3.2\times10^{-5}$ N/m | - |
| Bacterial mass density | 1100 $kg/m^3$<br>(*chosen slightly bigger than that of water*) | - |
| Maximum length of a pilus | 5 µm | (3) |
| Number of pili | 1-5 | (3) |
| Pilus angle variation | Standard deviation of pili angle distribution = 0-90 (deg.) | (4) |
| Pili spring constant | $2\times10^{-5}$ N/m | |
| Pili stall force | 100 pN | (5) |
| Pili retraction velocity | 1 µm/s | (6, 7) |
| Pili elongation velocity | 1 µm/s | (6) |
| Pili attachment probability | 0.8 (*the range in (2) is 0.1-0.3, but we have used an increased value so that we can obtain a reasonable twitching velocity*) | (2) |
| Pili detachment time, τ | 4 s | (8, 9) |
| Number of cells | 1-4 | - |

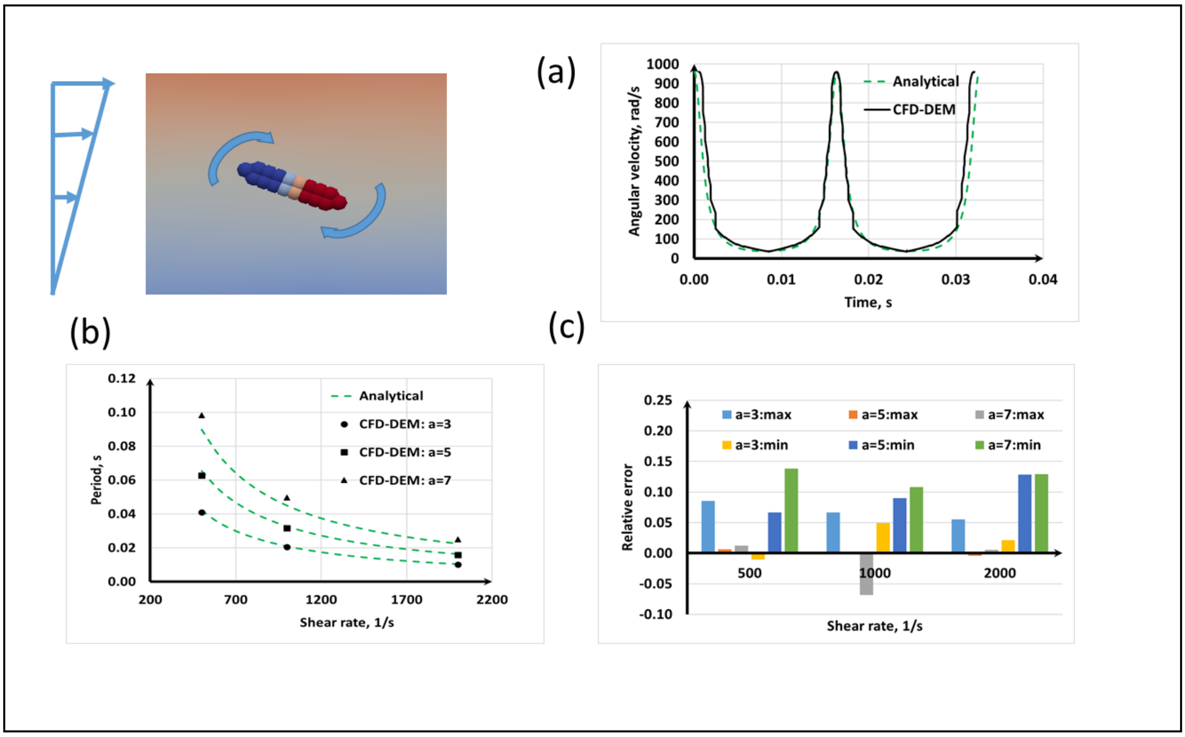


**Figure S1.** Model validation with Jeffery orbit of a rod shape bacteria: **(a)** Transient of the angular velocity at $a = 3$ and $\dot{\gamma} = 1000\ s^{-1}$; **(b)** period of the orbit; **(c)** relative error between analytical and CFD-DEM results for the maximum and minimum of the angular velocity at each case.

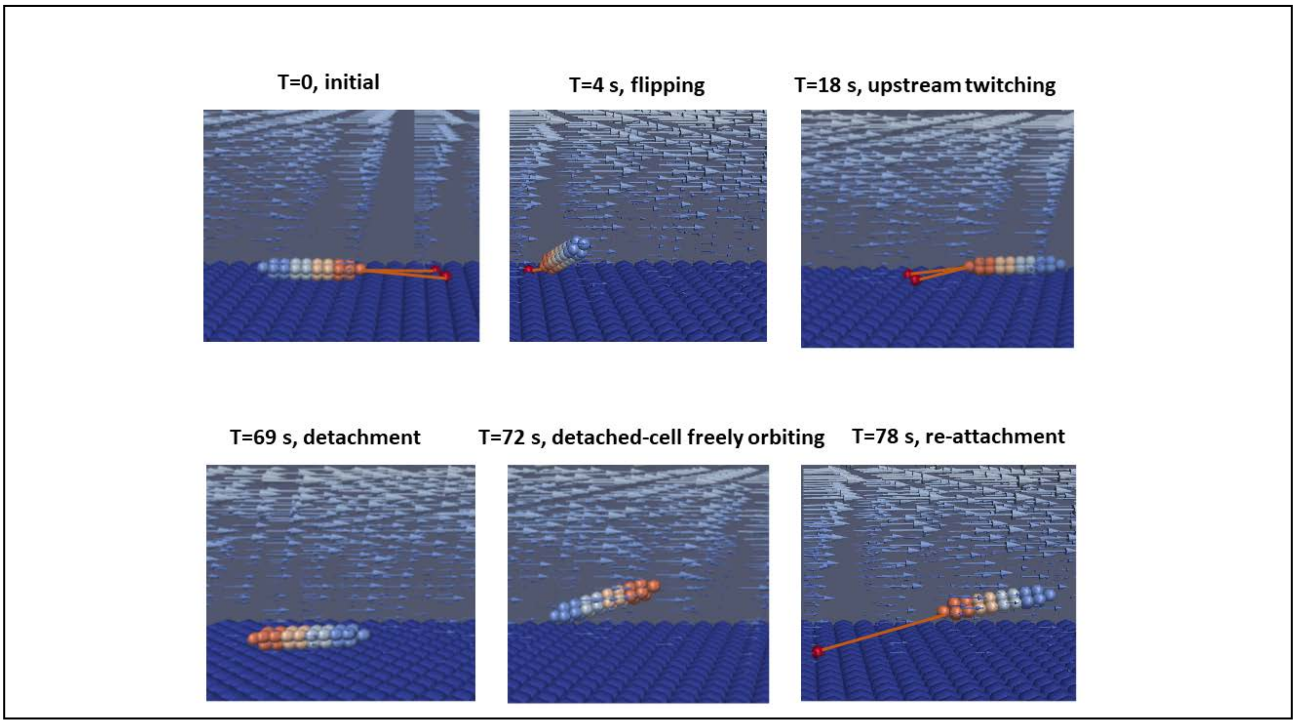


**Figure S2.** Bacterial twitching on a flat surface under shear flows.

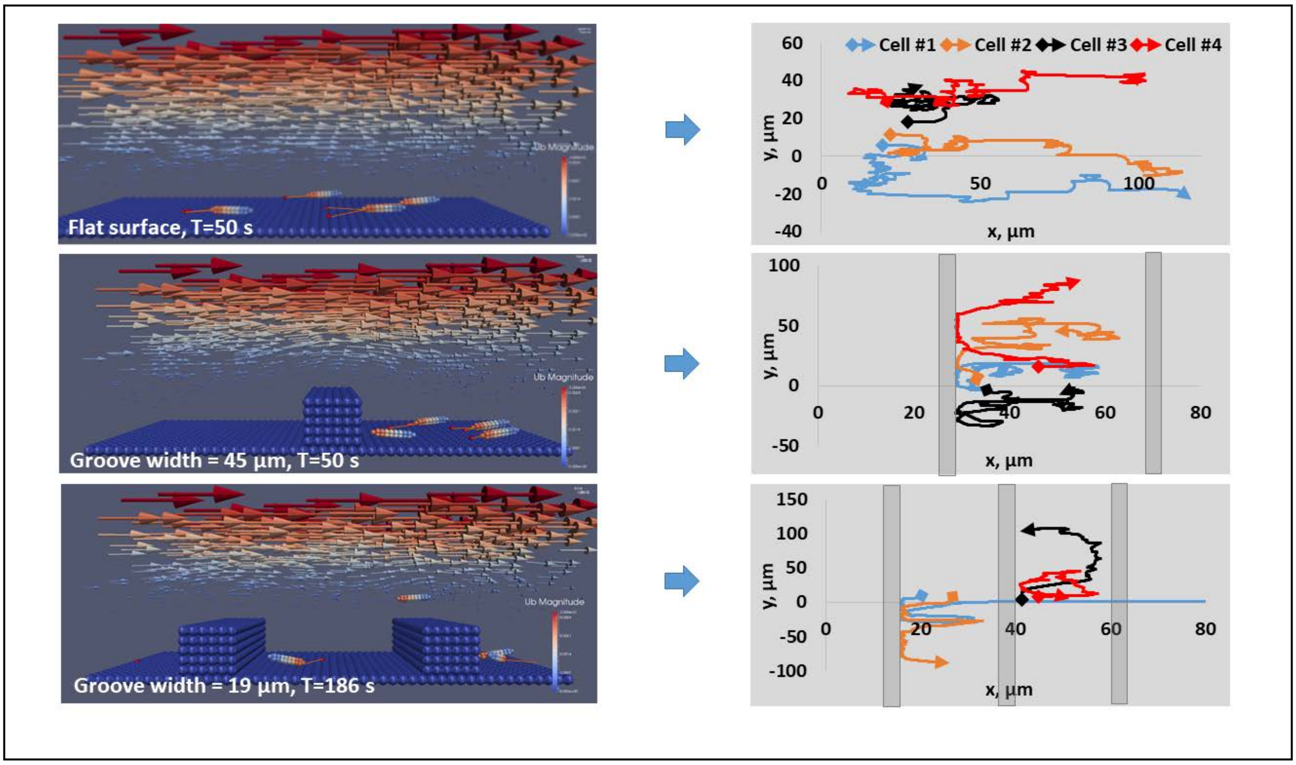


**Figure S3.** Bacterial twitching in shear flows under different groove width.
